# Haplotype-level metabarcoding of freshwater macroinvertebrate species: a prospective tool for population genetic analysis

**DOI:** 10.1101/2021.09.17.460774

**Authors:** Joeselle M. Serrana, Kozo Watanabe

## Abstract

The development and evaluation of DNA metabarcoding protocols for haplotype-level resolution require attention, specifically for population genetic analysis, i.e., parallel estimation of genetic diversity and dispersal patterns among multiple species present in a bulk sample. Further exploration and assessment of the laboratory and bioinformatics strategies are warranted to unlock the potential of metabarcoding-inferred population genetic analysis. Here, we assessed the inference of freshwater macroinvertebrate haplotypes from DNA metabarcoding data using mock samples with known Sanger-sequenced haplotypes. We also examined the influence of different DNA template concentrations and PCR cycles on detecting true haplotypes and the reduction of spurious haplotypes obtained from DNA metabarcoding. We tested our haplotyping strategy on a mock sample containing 20 specimens from four species with known haplotypes based on the 658-bp Folmer region of the mitochondrial cytochrome c oxidase gene. The read processing and denoising step resulted in 14 zero-radius operational taxonomic units (ZOTUs) of 421-bp length, with 12 ZOTUs having 100% match with 12 of the Sanger haplotype sequences. Quality passing reads relatively increased with increasing PCR cycles, and the relative abundance of each ZOTUs was consistent for each cycle number. This suggests that increasing the cycle number from 24 to 64 did not affect the relative abundance of quality passing filter reads of each ZOTUs. Our study demonstrated the ability of DNA metabarcoding to infer intraspecific variability while highlighting the challenges that need to be addressed before its possible applications to population genetic studies.

## INTRODUCTION

High throughput sequencing (HTS) of amplified markers, i.e., DNA metabarcoding, enables the simultaneous multi-species identification of community samples containing DNA from different origins (Taberlet et al., 2012). The advancements in sequencing platform technologies made it possible to generate millions of sequences from bulk (i.e., mixed, mass-collected, or community) samples and potentially identify most taxa, including the rare and inconspicuous ones (Creer et al., 2016). The number of sequences varies depending on the specificity of the barcoding gene and the species diversity present in the environment (Yu et al., 2012; Zinger et al., 2019).

The development and application of DNA metabarcoding for biodiversity surveys have been considered a game-changer for ecological research (Creer et al., 2016). The ability of the method to identify freshwater macroinvertebrates better than morphological identification has been demonstrated in previous literature (e.g., Hajibabaei et al., 2012; Serrana et al., 2019). However, other than the characterization of the communities’ taxonomic composition, DNA metabarcoding data can be used to infer biodiversity indices and is by now established as a robust method for environmental impact assessments of freshwater ecosystems (e.g., Aylagas et al., 2016; Deiner et al., 2017; Pawlowski et al., 2018; Serrana et al., 2018). For these applications, biodiversity is generally quantified with taxonomic, functional, or phylogenetic diversity at the community level (interspecific diversity) (Gotelli et al., 2017). However, intraspecific diversity or the genetic variation within species from community samples is not typically explored.

The inclusion of intraspecific diversity assessment would be beneficial in ecological monitoring and management strategies given that haplotype data are direct proxies for the spatio-temporal dynamics of populations and can substantially differ compared to community-level assessments (Taberlet et al., 2012). In particular, the assessment of changes in population size in response to environmental stressors (e.g., Monaghan et al., 2005; Weiss & Leese, 2016) are key areas of basic and applied ecological research (Sutherland et al., 2013). Quantifying gene flow between populations can examine the magnitude and mechanisms of population connectivity.

Previous literature has proposed using DNA metabarcoding for population genetic analysis (e.g., Bohmann et al., 2014; Adams et al., 2019; Arribas et al., 2020; Shum & Palumbi, 2021). Recent studies in freshwater environments were able to infer haplotypes from DNA metabarcoding of individually-tagged specimens (Shokralla et al., 2014), bulk macroinvertebrate samples (Elbrecht & Leese, 2015; Pedro et al., 2017; Zizka, Weiss & Leese, 2020), and from environmental DNA collected from water samples (Stat et al., 2017; Alam et al., 2020; Tsuji et al., 2020a; Tsuji et al., 2020b; Dugal et al., 2021), all of which highlighted the possibility of extracting haplotype or sequence variant information from HTS marker-gene sequencing data. DNA metabarcoding can be a faster and cost-effective alternative to the conventional approach that repeats genetic analysis for multiple individuals from different populations. Metabarcoding datasets may allow the comparison of population genetic structures of multiple species with different life stages and dispersal modes or abilities (Shum & Palumbi, 2021).

However, the application of DNA metabarcoding to population genetic analysis are highly influenced by the high number of reads containing sequencing errors, which could have occurred at different steps in the metabarcoding procedure, from polymerase chain reaction (PCR) amplification bias and errors to bioinformatics pipelines (Elbrecht et al., 2018; Turon et al., 2019). DNA template concentration and the number of PCR cycles introduce PCR bias and errors, mainly when applied to bulk samples (Pawluczyk et al., 2015; Dopheide et al., 2017; Collins et al., 2019). This makes it difficult to distinguish true haplotypes from erroneous sequencing noise (Turon et al., 2019). Thus, further exploration and assessment of the laboratory and bioinformatics strategies are required to unlock the potential of metabarcoding-based inference of haplotype information.

Here, we inferred haplotype information of freshwater macroinvertebrate species from DNA metabarcoding data. Using a mixed-species mock sample with known Sanger-sequenced haplotypes based on the 658-bp barcode region of the mitochondrial cytochrome oxidase I (mtCOI) gene, we tested whether the same COI haplotypes will be identified from the DNA metabarcoding data. We also examined the influence of varying DNA template concentration and PCR cycles on detecting true haplotypes and reducing spurious haplotypes obtained from DNA metabarcoding using the denoising parameters of unoise3 (Edgar, 2016). As DNA metabarcoding becomes more established and laboratory protocols and bioinformatics pipelines are continuously being developed, this study aimed to demonstrate that the method could be used to infer intraspecific genetic variability, showing promise for possible applications in population genetic analysis.

## MATERIALS AND METHODS

### Study species and Sanger-sequence haplotypes

We tested our haplotyping strategy on a mock sample by pooling extracted DNA of the 20 specimens from four species with known haplotypes assessed from published population genetics studies (*Amphinemura decemseta*; Gamboa et al., 2019) or our current DNA barcoding projects (*Kamimuria tibialis, Eucapnopsis bulba*, and *Epeorus latifolium*) (**Table 1**). Twenty haplotypes were identified based on the Sanger sequence of the 658-bp Folmer region of the mitochondrial cytochrome c oxidase I (mtCOI) gene using DnaSP 6.0 (Rozas et al., 2017).

**Table 1.** Sample and haplotype information table.

| Order | Family | Species | GenBank Accession | Sample Code | Conc. ng/uL<sup>†</sup> | Haplotype Information |  |  |  |  |  |
| --- | --- | --- | --- | --- | --- | --- | --- | --- | --- | --- | --- |
|  |  |  |  |  |  | 658-bp<sup>‡</sup> |  |  | 421-bp<sup>§</sup> |  |  |
|  |  |  |  |  |  | *h* | *Hd* | *Eta* | *h* | *Hd* | *Eta* |
| Plecoptera | Nemouridae | *Amphinemura decemseta* | MK132341 | AD1 | 0.01 |  |  |  |  |  |  |
|  |  |  | MK132323 | AD2 | 0.05 | 3 | 1.00 | 6 | 2 | 0.67 | 5 |
|  |  |  | MK132340 | AD3 | 0.10 |  |  |  |  |  |  |
|  | Perlidae | *Kamimuria tibialis* | MZ543957 | KT1 | 0.01 |  |  |  |  |  |  |
|  |  |  | MZ543958 | KT2 | 0.05 |  |  |  |  |  |  |
|  |  |  | MZ543959 | KT3 | 0.10 |  |  |  |  |  |  |
|  |  |  | MZ543960 | KT4 | 0.50 | 7 | 1.00 | 245 | 5 | 0.86 | 169 |
|  |  |  | MZ543961 | KT5 | 1.00 |  |  |  |  |  |  |
|  |  |  | MZ543962 | KT6 | 5.00 |  |  |  |  |  |  |
|  |  |  | MZ543963 | KT7 | 10.00 |  |  |  |  |  |  |
|  | Capniidae | *Eucapnopsis bulba* | Unr. Seq. | EB1 | 0.01 |  |  |  |  |  |  |
|  |  |  | Unr. Seq. | EB2 | 0.05 |  |  |  |  |  |  |
|  |  |  | Unr. Seq. | EB3 | 0.10 | 5 | 1.00 | 104 | 5 | 1.00 | 71 |
|  |  |  | Unr. Seq. | EB4 | 0.50 |  |  |  |  |  |  |
|  |  |  | Unr. Seq. | EB5 | 1.00 |  |  |  |  |  |  |
| Ephemeroptera | Heptageniidae | *Epeorus latifolium* | MZ543952 | EL1 | 0.01 |  |  |  |  |  |  |
|  |  |  | MZ543953 | EL2 | 0.05 |  |  |  |  |  |  |
|  |  |  | MZ543954 | EL3 | 0.10 | 5 | 1.00 | 19 | 5 | 1.00 | 14 |
|  |  |  | MZ543955 | EL4 | 0.50 |  |  |  |  |  |  |
|  |  |  | MZ543956 | EL5 | 1.00 |  |  |  |  |  |  |

Notes: <sup>†</sup>DNA concentration in the mock sample; <sup>‡</sup>Haplotypes based on the 658-bp Folmer region of the mtCOI gene; <sup>§</sup>421-bp internal COI fragment of the BF2 + BR2 primers. "*h*" refers to the number of haplotypes; "*Hd*" haplotype diversity; "*Eta*" Total number of mutations. "Unr. Seq." are unreleased sequences (Fasta files are provided in the supplementary materials).

The genealogical relationship of the haplotypes of each species via median-joining network was determined using a haplotype network tree (**Figure 1a)** drawn using the PopART software (Full-Feature Software for Haplotype Network Construction) (Leigh & Bryant, 2015). To examine the influence of varying input DNA template concentration on the detection of true haplotypes from the mock sample, a concentration gradient from the extracted DNA of the haplotypes from each species was prepared, i.e., 0.01, 0.05, 0.1, 0.5, and 1.0 ng/μL referred to as haplotypes 1 to 5, respectively (**Table 1**). Additional two samples with different concentrations of 5.0 and 10.0 ng/μL for *Kamimuria tibialis* were included, referred to as haplotype 6 and 7 (**Table 1**). DNA concentration was quantified with the QuantiFluor dsDNA system (Promega, Madison, WI, USA) on the Quantus Fluorometer (Promega, Madison, WI, USA).

**Figure 1.**
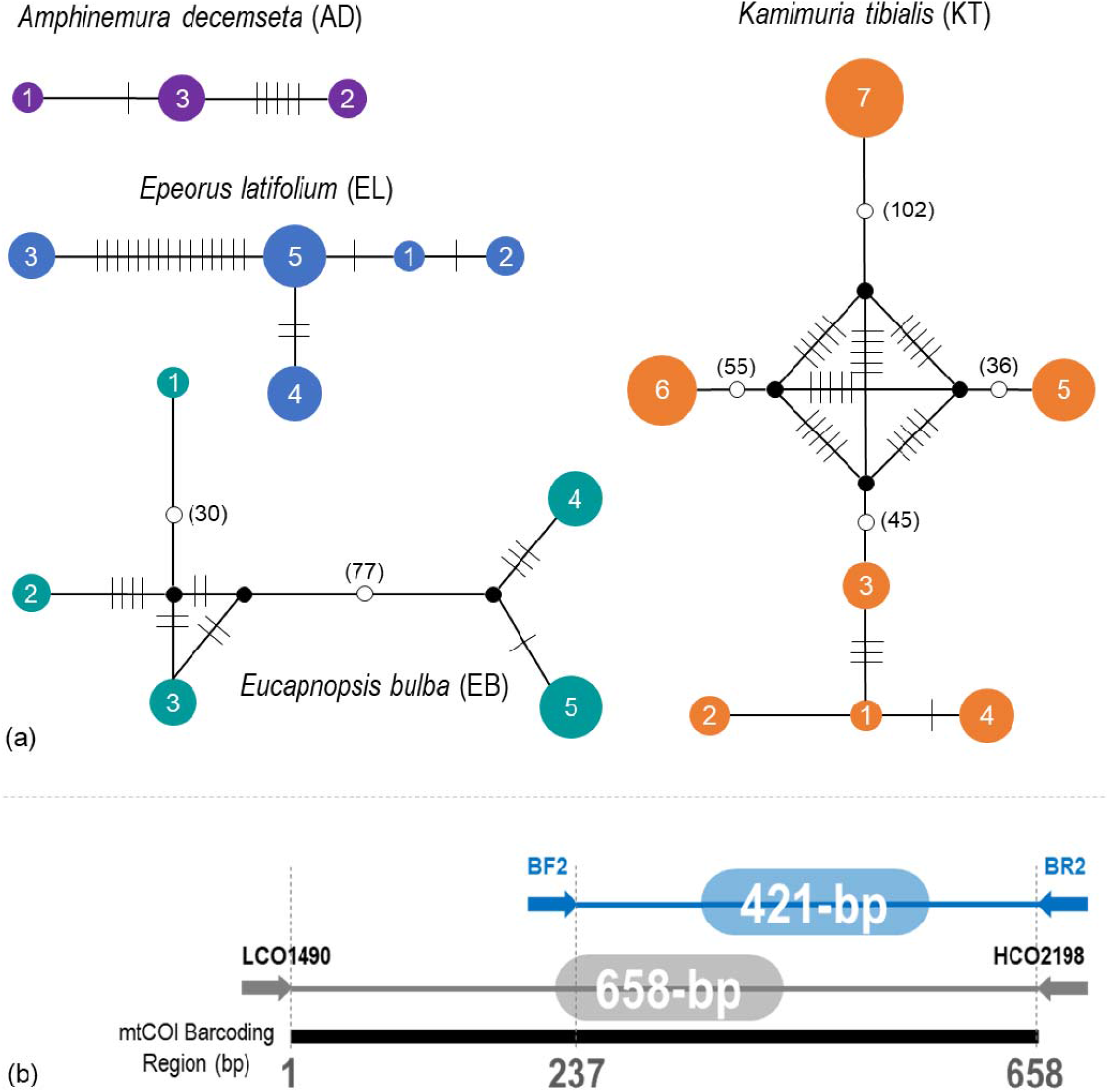
Median-joining haplotype networks showing the level of intraspecific variation within each of the four species based on the 658-bp mtCOI Sanger sequence (a). The number of mutations is represented as hatch marks or numbers. Position of the sequenced amplicon (BF2 & BR2) along the mtCOI barcode region (b).

### Library preparation and next-generation sequencing

Fifteen μl each of the 20 haplotype DNA extracts adjusted to their respective concentrations were pooled as one mock sample. Elbrecht and Steinke’s (2019) fusion primers (BF2 and BR2; **Figure 1b**) for freshwater arthropods were used to construct amplicon libraries following a one-step polymerase chain reaction (PCR) protocol. The PCR master mix consists of 0.25 μl Phusion, 0.75 μl DMSO, one μl dNTPs, 1.25 μl each of the fusion forward and reverse primers (10 μM), five μl HF Buffer (New England Biolabs), and 15.5 μl of PCR-grade water. PCR cycling conditions were 30 seconds of initial denaturation at 98°C, followed by 20 cycles of 10 seconds denaturation at 98°C, 30 seconds annealing at 55°C, 30 seconds extension at 72 °C, and a final extension step of 5 minutes at 72°C. The same PCR protocol was done for eleven other cycles from 24 to 64 (increments of 4) with triplicate PCR amplification, including negative controls for each PCR cycle (also in triplicates).

The resulting 72 PCR amplicons were pooled according to replicates and size selected via solid-phase reversible immobilization (SPRI) beads. Each replicate was quantified via qPCR with the KAPA Library Quantification Kit (Kapa Biosystems, Wilmington, MA, USA). Before sequencing, quality was assessed via the DNA 1000 assay using the Agilent 2100 BioAnlyzer (Agilent Technologies, Palo Alto, CA, USA). The replicate amplicon libraries were normalized to 2nM before pooling to ensure even read output distribution between replicates. Paired-end sequencing of the 2nM pooled library spiked with 20% PhiX was performed in paired-end 300 cycles sequencing on the Illumina MiSeq system.

### Sequence processing and haplotype inference

The raw Illumina MiSeq paired-end reads were demultiplexed according to sample tags via the R package JAMP v.0.67 (http://github.com/VascoElbrecht/JAMP) (Elbrecht et al., 2018) and were quality-checked with FastQC (https://www.bioinformatics.babraham.ac.uk/projects/fastqc/). We demultiplexed 14.7 million reads assigned to each of the 72 samples. The paired-end reads were then merged via the JAMP pipeline, primer stripped, truncated at 421-bp, quality filtered with a maximum expected error filtering value of 2, and dereplicated excluding singletons using the USEARCH v11.0.667 pipeline (Edgar, 2013). To extract individual haplotypes from the dereplicated sequences, we employed a denoising strategy using unoise3 (Edgar, 2016). The zero-radius operational taxonomic unit (ZOTUs) sequences were mapped against the 658-bp mtCOI Sanger sequences of the mock samples using the UPARSE-REF algorithm (Edgar, 2013). The command is designed for validating mock community sequencing experiments where the set of biological sequences in the sample is known. Note that amplicon sequence variants (ASVs), exact sequence variants (ESVs), sub-OTUs, OTUs with 100% sequence similarity, and ZOTUs are synonymous (Porter & Hajibabaei, 2018), but we used the latter term in this study since we employed unoise3’s algorithm (Edgar, 2016). We also performed phylogenetic inference via neighbor-joining analysis of the Sanger and the ZOTU sequences employing the Jukes-Cantor substitution model with bootstrapping (1000) via the online MAFFT multiple sequence alignment software version 7 (https://mafft.cbrc.jp/alignment/server/large.html) (Katoh, Rozewicki & Yamada, 2019). Prior to the analysis, the 658-bp Sanger sequences of the mock samples were truncated to the 421-bp length of the BF2 and BR2 barcode.

Data visualization (i.e., boxplots and bubble plot) and statistical analysis were performed in R v.4.0.2 (R Core Team, 2019). The default Wilcoxon T-test analysis was performed to compare the read abundances of the raw, merged, and ZOTU-assigned sequences between the DNA template and negative controls for each cycle using the function stat_compare_means().

## RESULTS

### Sanger sequence haplotypes

The mock sample consisted of four different species, i.e., *Amphinemura decemseta* (Nemouridae), *Kamimuria tibialis* (Perlidae), *Eucapnopsis bulba* (Capniidae), and *Epeorus latifolium* (Heptageniidae) from two freshwater insect orders (Table 1). However, upon trimming the Sanger sequences into the 421-bp length of the BF2 and BR2 barcode region to complement with the amplified region in the DNA metabarcoding data, three of the *K. tibialis* Sanger haplotypes (i.e., KT1, KT3, and KT4) were grouped into one, similar to the two haplotypes of *A. decemseta* (i.e., AD1 and AD3). Hence, the 20 haplotypes from the 658-bp long fragment of the mtCOI barcoding region (Folmer et al., 1994) were reduced to 17 haplotypes after trimming to 421-bp length.

### Read abundance, reference sequence match, and phylogenetic inference

Except for the raw sequence and paired-end merging step at cycle 20, the read abundances of the DNA template samples and the negative controls were significantly different (**Figure 2a**). From the dereplicated reads, 462,665 were assigned into 14 ZOTUs (also referred to as “DNA metabarcoding haplotypes” in this study) (**Table S1**) of 421-bp length. For assessing the different PCR cycle numbers, all of the ZOTUs were detected from cycles 24 to 64. For cycle 20, only 8 of the 14 ZOTUs were represented, all of which match haplotypes with high concentrations (i.e., KT7, KT6, KT5, EL5, EL4, EB5, EB4, and EB3) (**Figure 2b**). Quality passing reads relatively increased with increasing cycle numbers, and the relative abundance of each ZOTUs was relatively consistent across the cycle numbers (**Table S1**). Notably, four ZOTUs were detected in the negative samples from different cycles. ZOTU04 was present on nine negative samples from cycles 28 to 64, ZOTU09 and ZOTU11 on two negative samples, and ZOTU08 and ZOTU10 on one, i.e., cycle 60 and 64, respectively. However, it should be noted that most of these occurrences in the negative samples were singleton or doubleton reads, with one sample (i.e., 40BR1) having the highest detection of only 22 reads.

**Figure 2.**
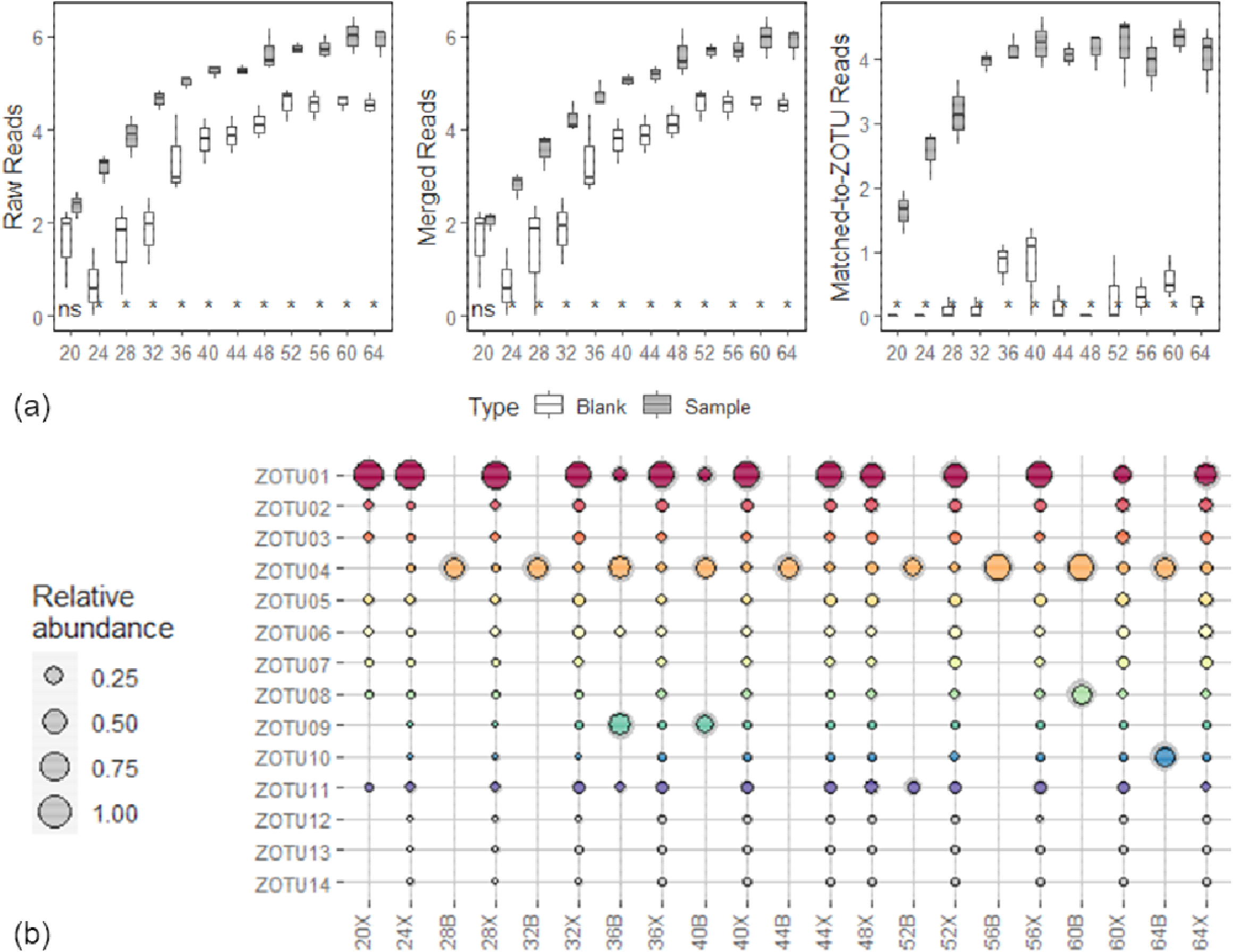
Comparison of read abundance (log-transformed) at different read processing steps per cycle and between the DNA template samples and the negative controls (a). Significance code: ‘**’ associated with a variable at *p* < 0.01, and ‘*’ at *p* < 0.05. Bubble plot showing the relative abundance of ZOTUs per PCR cycle (b). Circle size represents the Mean values of three replicates, and the gray shadow as standard deviation (SD). “-X” are samples with DNA template; “-B” are negative controls.

After mapping the ZOTU sequences against the 20 Sanger sequence-haplotypes with 658-bp, 12 ZOTUs comprised of 450,793 (97%) matched-to-ZOTU reads had 100% sequence match against 12 of the Sanger haplotypes of the mock reference sequence (**Table S2**). The remaining eight Sanger haplotypes that were not detected from the DNA metabarcoding dataset were all the *A. decemseta* samples (AB1: 0.01, AB2: 0.05, and AB3: 0.10 ng/μL), two *E. bulba* (EB1: 0.01, and EB2: 0.05 ng/μL), *E. latifolium* (EL1: 0.01 ng/μL), and two *K. tibialis* (KT1:0.01, and KT3: 0.10 ng/μL). Based on the neighbor-joining tree (**Figure 3**), the remaining two ZOTUs (i.e., ZOTU10 and ZOTU14) without a 100% taxonomic match against the Sanger sequences clustered with the *A. decemseta* sequences.

**Figure 3.**
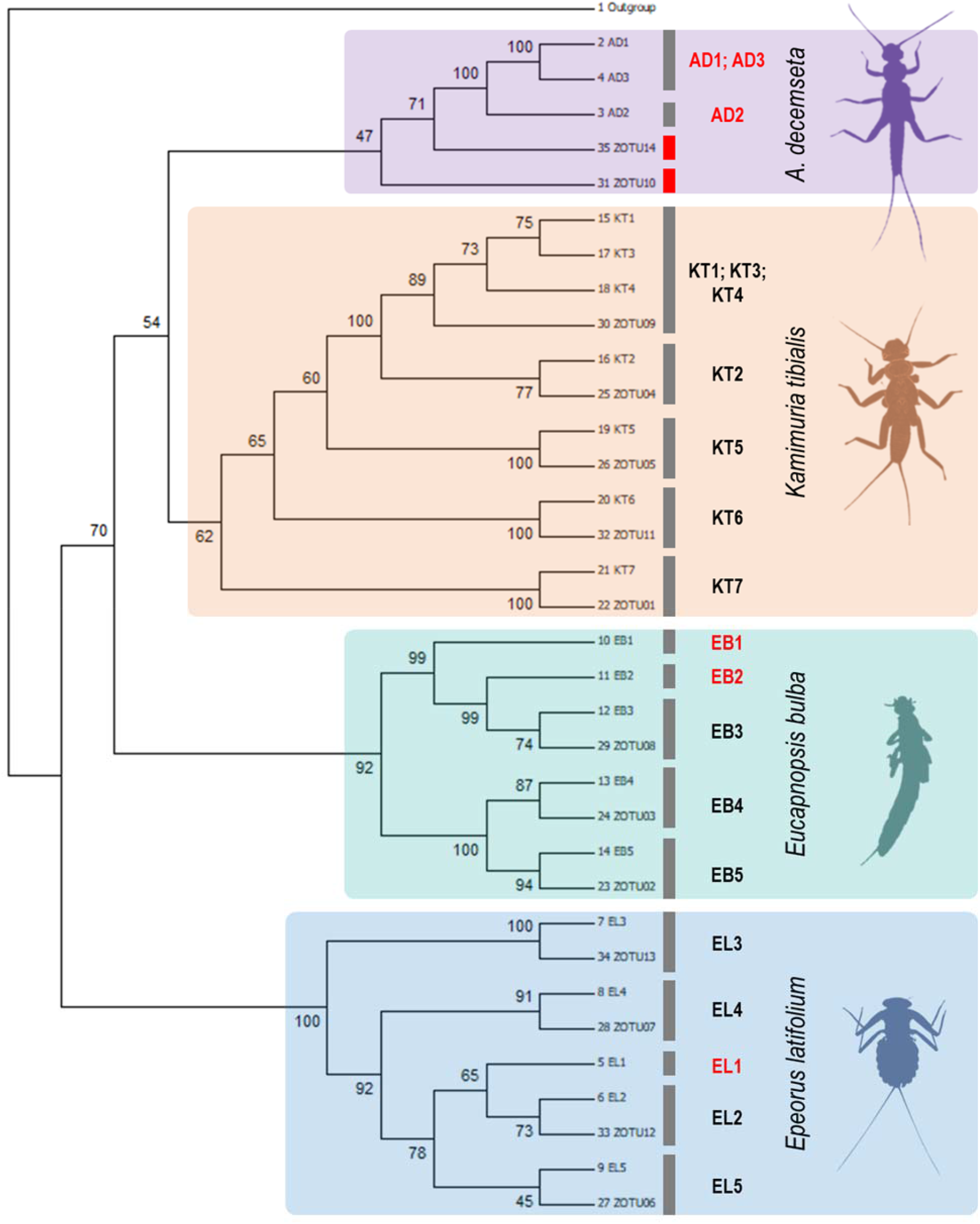
Neighbor-joining tree of the mock and the zero-radius operational taxonomic units (ZOTU) sequences. Sequences are highlighted based on species. The red bar represents a ZOTU without a taxonomic match, and the text in red represents a Sanger haplotype without a ZOTU match.

## DISCUSSION

Using a multi-species mock sample with known haplotypes based on Sanger sequencing of the highly variable mtCOI gene region, we presented the feasibility and limitations of using DNA metabarcoding data to extract intraspecific genetic diversity information by denoising the sequences into ZOTUs. Most recent studies inferred intraspecific diversity from macroinvertebrate DNA metabarcoding data were also based on denoising algorithms (e.g., Elbrecht et al., 2018; Turon et al., 2019; Brandt et al., 2021). Before denoising, our read processing steps, i.e., paired-end merging, stripping adapter and primer sequences, barcode length truncation, quality filtering, and dereplication, were performed mainly following default and stringent settings. It is also important to note that we truncated the sequence reads to the entire length of the BF2 and BR2 barcode (i.e., 421-bp) to prevent the generation of shorter-than-barcode ZOTUs. We also discarded singletons, i.e., ZOTUs containing only one sequence read.

The four species selected in the mock sample have known haplotypes assessed from a published population genetics study (*Amphinemura decemseta*; Gamboa et al., 2019) or our current DNA barcoding projects through the sequencing of the 658-bp region of the mtCOI gene. Upon truncating the mock sequences to the 421-bp length of the BF2 and BR2 barcode, two species, i.e., *A. decemseta* and *K. tibialis* were reduced from 3 to 2 and 7 to 5 haplotypes, respectively. This means that some of the polymorphic sites that distinguished the haplotypes from each other were discarded from shortening the 658-bp barcoding region to 421-bp. Twelve of the 14 ZOTUs inferred from the DNA metabarcoding sequences had perfect sequence matches against 12 of the 20 (658-bp) or 17 (421-bp) Sanger haplotypes. The *K. tibialis* haplotype with a relatively high input DNA template concentration (i.e., 0.10 ng/μL) was likely not a false negative detection. Its reads were present in the metabarcoding data but merged with the other *K. tibialis* haplotype.

We report that initial DNA template concentration significantly influenced the detection of individuals from a mock community sample. This observation is in accordance with previous studies which reported that samples or taxa with low DNA template concentrations had lower detection probability (Martins et al., 2020). Accordingly, abundant taxa or samples with high biomass tend to have higher detection probabilities than those rare, smaller, or have low biomass from mixed-community samples (Carew, Coleman & Hoffmann, 2018; Erdozain et al., 2019; Serrana et al., 2019). The difference in biomass affects haplotypes’ detection since most large specimens would be retained after read processing. These factors need to be addressed when metabarcoding-based haplotyping is used to infer abundance-based analysis for population genetics applications.

On the other hand, all of the ZOTUs were detected from cycles 24 to 64. For cycle 20, only 8 of the 14 ZOTUs were represented, all of which match haplotypes with high concentrations. Quality passing reads relatively increased with increasing cycle numbers, and the relative abundance of each ZOTUs was consistent for each cycle number. This suggests that increasing the PCR cycle from 24 to 64 did not affect the relative abundance of quality passing filter reads of each ZOTUs. Our findings align with previous studies that reported small or no effect of PCR cycle number on amplification bias (Sipos et al., 2007; Krehenwinkel et al., 2017). This contrasts with other reports that increasing PCR cycles reduces the proportion of sequences with low starting DNA or less well-amplified species in the mock sample (Piñol et al., 2015).

Moreover, higher PCR cycles have been reported to increase the formation of chimeric sequences and amplification bias (Aylagas et al., 2016); that is why some metabarcoding protocols discourage increasing the PCR cycles above 30. However, a literature survey on bulk metabarcoding studies presented that 73% of reports used more than 30 PCR cycles to circumvent primer annealing issues or to amplify samples with low amounts of DNA (van der Loos & Nijland, 2020). Our findings proved otherwise wherein low template samples were undetected or not amplified even after increasing the cycle number to 64. The relative abundances of the samples detected were relatively consistent for each cycle. However, we note that the sequence diversity in our mock sample was relatively low (i.e., four species with 20 haplotypes). A more diverse community might present a different pattern in PCR cycle effects, hence, warrants further evaluation.

The proper selection of the DNA marker in DNA metabarcoding assay is crucial because all of the downstream analyses, e.g., species detection and identification, rely on the marker’s ability to amplify and discriminate the representative taxa of the target organisms (Piper et al., 2019). The mtCOI gene has a relatively high mutation rate and is capable of detecting intraspecific variation. Thus, its widespread use for population genetics studies (e.g., Hajibabaei et al., 2007; Curry et al., 2018) and has been widely used for DNA metabarcoding macroinvertebrates to date. With this, intraspecific variation can be extracted from community-based samples using various algorithms for sequence clustering and phylogenetic rates (Giusti et al., 2017; Vamos, Elbrecht & Leese, 2017; Elbrecht et al., 2019). However, many HTS platforms to date still have strict limitations in sequence length, not utilizing the full-length of the mtCOI barcode (Piper et al., 2019).

As we have observed in this study, some species lost nucleotide information to distinguish the haplotypes from each other due to shortening the barcode region. Hence, the inability of shorter marker regions to differentiate haplotypes of certain species requires further development and evaluation. On the other hand, the advancements in sequencing technologies to generate longer sequences could allow the full-length generation of longer DNA barcodes from community samples (Baloğlu et al., 2021), resolving the limitations of short amplicon sequencing in DNA metabarcoding. Longer reads would allow the generation of high-resolution mitochondrial haplotype data, with potential applications for demographic history and selection analyses (Sigsgaard et al., 2019). However, given the high raw read error rate from these long-read sequencing platforms that range from 10-22% (i.e., the Oxford Nanopore MinION™), further assessment and exploration of library preparation and error-correction methods are recommended (Baloğlu et al., 2021) for its utilization for DNA metabarcoding studies, more so for haplotype-level inference.

The two DNA metabarcoding haplotypes (i.e., ZOTU10 and ZOTU14) that failed to have a perfect sequence match against the Sanger sequences clustered with the *A. decemseta* samples. Although we could identify them as *A. decemseta* sequences based on the phylogenetic approach, metabarcoding failed to detect a 100% match of the Sanger haplotypes of this species from the mock sample. We could not rule out PCR amplification, primer bias, or sequencing errors as the reason for these false positive detections or spurious haplotypes obtained from the metabarcoding data, even if we performed relatively strict read quality filtering parameters. Elbrecht et al. (2018) reported that artificial or false haplotypes could not be fully excluded even with stringent filtering settings due to undetected chimeric sequences or systematic sequencing errors that might persist across replicates.

Additionally, nine blank samples had sequences for ZOTU04, ZOTU08, ZOTU09, ZOTU11, and ZOTU10. Although most of these occurrences were singleton or doubleton reads, the presence of these reads in the negative controls might be due to tag-jumps or the amplification of sequences carrying false combinations of used nucleotide tags that is common for dual indexed libraries (Esling, Lejzerowicz & Pawlowski, 2015; Schnell, Bohmann & Gilbert, 2015; Zizka et al., 2019). Hence, the challenges with tag jumping and contamination between libraries require attention to alleviate false read-to-sample assignments that would be problematic once the method is employed to environmental samples.

## CONCLUSION

We demonstrated that haplotype information could be extracted from mixed community samples of benthic macroinvertebrates. We recovered 14 zero-radius operational taxonomic units (ZOTUs) of 421-bp length, with 12 ZOTUs having 100% match with 12 of the mock haplotype sequences. Moreover, quality passing reads relatively increased with increasing PCR cycle numbers. The relative abundance of each ZOTUs was consistent across the cycle numbers suggesting that increasing the cycles did not affect the relative abundance of quality passing filter reads. As DNA metabarcoding becomes more established and laboratory protocols and bioinformatics pipelines are continuously being developed, our study demonstrated that the method could be used to infer intraspecific variability, showing promise for possible applications and highlighting the challenges that need to be addressed before haplotype-level metabarcoding can entirely be used for population genetic studies.

## Acknowledgments

This work was supported by the Japan Society for the Promotion of Science (JSPS) Grant-in-Aid for Scientific Research (Grant No. 19K21996 and 19H02276). We thank Dr. Dávid Murányi, Dr. Maribet Gamboa and Dr. Sakiko Yaegashi for the EPT samples and their molecular data. We also thank Dr. Naohito Tokunaga of the Division of Analytical Bio-Medicine for his assistance in performing high-throughput sequencing on the Illumina MiSeq platform of the Advanced Research Support Center (ADRES), Ehime University.

## Author contributions

K.W. and J.M.S. conceptualized and designed the study; J.M.S. performed laboratory work and bioinformatics analyses. J.M.S. and K.W. interpreted the data and wrote the manuscript.

## Conflict of interest statement

The authors declare that they have no known competing financial interests or personal relationships that could have influenced the work reported in this paper.

## Data availability statement

The raw sequence data were deposited into the National Center for Biotechnology Information (NCBI) Sequence Read Archive (SRR14631855). The Sanger (GenBank accession numbers also listed in Table 1) and ZOTU sequences (Fasta format), other input data, and the R markdown document implementing the analyses contained in this manuscript are available in the figshare data repository at https://doi.org/10.6084/m9.figshare.15090180.v1 (Serrana & Watanabe, 2021).

## Supporting information

The supplementary data to this article is provided as Additional file 1 (DOCX).

## Notes

https://doi.org/10.6084/m9.figshare.15090180.v1

### Competing Interest Statement

The authors have declared no competing interest.

