## Supplementary Information for "Haplotype-level metabarcoding of freshwater macroinvertebrate species: a prospective tool for population genetic analysis"

*Running Title: DNA metabarcoding-based haplotyping*

### Authors and Affiliations

Joeselle M. Serrana and Kozo Watanabe

Center for Marine Environmental Studies, Ehime University, Bunkyo-cho 3, Matsuyama, Ehime 790-8577, Japan

### Correspondence

Prof. Kozo Watanabe, Ph.D.; Address: Molecular Ecology and Health (MEcoH) Laboratory, Engineering Building No. 2, Ehime University, Bunkyo-cho 3, Matsuyama, Ehime 790-8577, Japan;

#### Supplementary Tables

##### **Table S1** Absolute read abundance of the zero-radius operational taxonomic units (ZOTUs) of each PCR cycle assessed in the study.

| **Sample** | | **ZOTU01** | **ZOTU02** | **ZOTU03** | **ZOTU04** | **ZOTU05** | **ZOTU06** | **ZOTU07** | **ZOTU08** | **ZOTU09** | **ZOTU10** | **ZOTU11** | **ZOTU12** | **ZOTU13** | **ZOTU14** |
| --- | --- | --- | --- | --- | --- | --- | --- | --- | --- | --- | --- | --- | --- | --- | --- |
| PCR Cycles | 20XR1 | 61 | 5 | 4 | - | 7 | 3 | 3 | 1 | - | - | 3 | - | - | - |
|  | 20XR2 | 16 | 1 | 1 | - | - | - | - | - | - | - | - | - | - | - |
|  | 20XR3 | 45 | - | - | - | - | 1 | - | - | - | - | - | - | - | - |
|  | 24XR1 | 465 | 15 | 22 | 12 | 46 | 32 | 20 | 5 | 1 | 2 | 34 | 1 | 1 | 3 |
|  | 24XR2 | 115 | 2 | 1 | 1 | 3 | - | 2 | 2 | - | - | 1 | - | - | - |
|  | 24XR3 | 533 | 5 | 2 | 1 | 6 | 2 | 1 | 1 | 1 | - | 13 | - | - | - |
|  | 28XR1 | 2,724 | 276 | 229 | 152 | 368 | 348 | 186 | 62 | 18 | 32 | 294 | 24 | 26 | 13 |
|  | 28XR2 | 424 | 4 | 4 | - | 5 | 1 | 1 | 5 | - | - | 9 | - | - | - |
|  | 28XR3 | 1,248 | 9 | 7 | 2 | 5 | 3 | - | 5 | 1 | - | 42 | - | - | - |
|  | 32XR1 | 3,696 | 805 | 785 | 525 | 1,031 | 947 | 608 | 208 | 71 | - | 860 | 68 | 116 | 55 |
|  | 32XR2 | 3,764 | 392 | 242 | 135 | 606 | 319 | 210 | 79 | 17 | 20 | 286 | 16 | 27 | 16 |
|  | 32XR3 | 10,064 | 398 | 242 | 139 | 541 | 315 | 59 | 22 | 11 | 13 | 888 | 12 | 21 | 13 |
|  | 36XR1 | 6,184 | 2,343 | 2,255 | 1,986 | 2,252 | 2,485 | 1,795 | 1,222 | 488 | 642 | 1,550 | 532 | 658 | 374 |
|  | 36XR2 | 8,716 | 61 | 16 | 1 | 14 | 5 | - | 15 | - | - | 1,472 | - | - | 1 |
|  | 36XR3 | 9,420 | 391 | 206 | 4 | 20 | 13 | 21 | 165 | - | - | 349 | 1 | 5 | - |
|  | 40XR1 | 7,490 | 4,312 | 4,080 | 4,138 | 3,662 | 4,267 | 3,001 | 3,376 | 1,303 | 1,370 | 2,514 | 1,108 | 1,429 | 1,068 |
|  | 40XR2 | 11,781 | 1,377 | 562 | 57 | 735 | 229 | 424 | 350 | 6 | 6 | 2,396 | 42 | 96 | - |
|  | 40XR3 | 6,925 | - | - | - | - | 2 | - | 29 | - | - | 8 | - | - | - |
|  | 44XR1 | 3,231 | 2,544 | 1,842 | 1,759 | 2,287 | 2,366 | 1,465 | 573 | 217 | 448 | 612 | 228 | 393 | 219 |
|  | 44XR2 | 6,120 | 416 | 93 | 9 | 66 | 9 | 4 | 22 | - | 1 | 749 | - | 1 | - |
|  | 44XR3 | 10,612 | 40 | 64 | 1 | 34 | 6 | 4 | 111 | 1 | - | 448 | - | - | - |
|  | 48XR1 | 4,241 | 2,717 | 1,961 | 2,258 | 2,968 | 2,442 | 1,558 | 1,244 | 256 | 490 | 980 | 231 | 371 | 229 |
|  | 48XR2 | 5,052 | 21 | 23 | - | 1 | - | 5 | 16 | 1 | - | 1,151 | - | 1 | - |
|  | 48XR3 | 9,720 | 2,935 | 1,465 | 1,149 | 2,033 | 212 | 66 | 1,455 | 73 | 24 | 1,490 | 24 | 14 | 17 |
|  | *Cont.* |  |  |  |  |  |  |  |  |  |  |  |  |  |  |
|  | 52XR1 | 5,899 | 5,046 | 3,763 | 3,113 | 4,243 | 4,283 | 2,922 | 2,222 | 462 | 1,761 | 1,360 | 598 | 926 | 589 |
|  | 52XR2 | 10,055 | 3,284 | 2,571 | 1,150 | 4,287 | 3,351 | 2,374 | 532 | 112 | 297 | 2,132 | 130 | 301 | 110 |
|  | 52XR3 | 3,325 | - | 2 | - | 1 | 3 | - | 31 | 1 | - | 182 | - | 1 | - |
|  | 56XR1 | 4,114 | 2,702 | 2,166 | 2,000 | 2,605 | 2,551 | 1,794 | 1,384 | 289 | 440 | 1,005 | 225 | 392 | 286 |
|  | 56XR2 | 7,480 | 241 | 113 | - | 242 | 7 | 7 | 47 | 2 | 1 | 1,815 | - | 1 | 1 |
|  | 56XR3 | 2,652 | 183 | 69 | 2 | 8 | 2 | 2 | 105 | 1 | - | 54 | - | - | - |
|  | 60XR1 | 4,363 | 2,823 | 2,666 | 1,850 | 2,867 | 2,424 | 1,566 | 888 | 185 | 449 | 946 | 156 | 324 | 169 |
|  | 60XR2 | 8,085 | 4,792 | 4,354 | 3,508 | 2,623 | 4,413 | 2,799 | 1,669 | 488 | 1,130 | 3,196 | 497 | 795 | 462 |
|  | 60XR3 | 4,990 | 700 | 1,720 | 860 | 2,383 | 267 | 188 | 553 | 76 | 40 | 418 | 7 | 19 | 15 |
|  | 64XR1 | 3,297 | 1,927 | 1,557 | 1,229 | 1,983 | 1,935 | 1,262 | 563 | 140 | 380 | 615 | 138 | 202 | 87 |
|  | 64XR2 | 5,110 | 3,796 | 2,983 | 2,517 | 3,558 | 5,041 | 2,680 | 998 | 277 | 418 | 707 | 190 | 444 | 179 |
|  | 64XR3 | 2,357 | 270 | 85 | 4 | 15 | 8 | 13 | 37 | 2 | 1 | 76 | - | 6 | - |
| Negative Control | 20BR1 | - | - | - | - | - | - | - | - | - | - | - | - | - | - |
|  | 20BR2 | - | - | - | - | - | - | - | - | - | - | - | - | - | - |
|  | 20BR3 | - | - | - | - | - | - | - | - | - | - | - | - | - | - |
|  | 24BR1 | - | - | - | - | - | - | - | - | - | - | - | - | - | - |
|  | 24BR2 | - | - | - | - | - | - | - | - | - | - | - | - | - | - |
|  | 24BR3 | - | - | - | - | - | - | - | - | - | - | - | - | - | - |
|  | 28BR1 | - | - | - | - | - | - | - | - | - | - | - | - | - | - |
|  | 28BR2 | - | - | - | 1 | - | - | - | - | - | - | - | - | - | - |
|  | 28BR3 | - | - | - | - | - | - | - | - | - | - | - | - | - | - |
|  | 32BR1 | - | - | - | - | - | - | - | - | - | - | - | - | - | - |
|  | 32BR2 | - | - | - | 1 | - | - | - | - | - | - | - | - | - | - |
|  | 32BR3 | - | - | - | - | - | - | - | - | - | - | - | - | - | - |
|  | 36BR1 | 2 | - | - | 5 | - | - | - | - | - | - | - | - | - | - |
|  | 36BR2 | - | - | - | 1 | - | - | - | - | 1 | - | - | - | - | - |
|  | 36BR3 | - | - | - | 1 | - | 1 | - | - | 9 | - | 1 | - | - | - |
|  | 40BR1 | 8 | - | - | 13 | - | - | - | - | 1 | - | - | - | - | - |
|  | 40BR2 | - | - | - | 4 | - | - | - | - | 7 | - | - | - | - | - |
|  | 40BR3 | - | - | - | - | - | - | - | - | - | - | - | - | - | - |
|  | *Cont.* |  |  |  |  |  |  |  |  |  |  |  |  |  |  |
|  | 44BR1 | - | - | - | - | - | - | - | - | - | - | - | - | - | - |
|  | 44BR2 | - | - | - | 2 | - | - | - | - | - | - | - | - | - | - |
|  | 44BR3 | - | - | - | - | - | - | - | - | - | - | - | - | - | - |
|  | 48BR1 | - | - | - | - | - | - | - | - | - | - | - | - | - | - |
|  | 48BR2 | - | - | - | - | - | - | - | - | - | - | - | - | - | - |
|  | 48BR3 | - | - | - | - | - | - | - | - | - | - | - | - | - | - |
|  | 52BR1 | - | - | - | - | - | - | - | - | - | - | - | - | - | - |
|  | 52BR2 | - | - | - | 6 | - | - | - | - | - | - | 2 | - | - | - |
|  | 52BR3 | - | - | - | - | - | - | - | - | - | - | - | - | - | - |
|  | 56BR1 | - | - | - | 1 | - | - | - | - | - | - | - | - | - | - |
|  | 56BR2 | - | - | - | 3 | - | - | - | - | - | - | - | - | - | - |
|  | 56BR3 | - | - | - | - | - | - | - | - | - | - | - | - | - | - |
|  | 60BR1 | - | - | - | 2 | - | - | - | - | - | - | - | - | - | - |
|  | 60BR2 | - | - | - | 1 | - | - | - | - | - | - | - | - | - | - |
|  | 60BR3 | - | - | - | - | - | - | - | 8 | - | - | - | - | - | - |
|  | 64BR1 | - | - | - | - | - | - | - | - | - | - | - | - | - | - |
|  | 64BR2 | - | - | - | 1 | - | - | - | - | - | - | - | - | - | - |
|  | 64BR3 | - | - | - | - | - | - | - | - | - | 1 | - | - | - | - |
| **Total** | | **174,374** | **44,833** | **36,155** | **28,562** | **41,505** | **38,292** | **25,040** | **17,997** | **4,500** | **7,965** | **28,655** | **4,228** | **6,570** | **3,906** |

##### **Table S2** Taxonomic match of the zero-radius operational taxonomic units (ZOTUs) from the DNA metabarcoding sequences against the 658-bp mtCOI Sanger sequences.

| **ZOTU** | **Order** | **Family** | **Genus** | **Species** | **Sanger Haplotype** |
| --- | --- | --- | --- | --- | --- |
| ZOTU01 | Plecoptera | Perlidae | *Kamimuria* | *Kamimuria tibialis* | KT7 |
| ZOTU02 | Plecoptera | Capniidae | *Eocapnosis* | *Eucapnopsis bulba* | EB5 |
| ZOTU03 | Plecoptera | Capniidae | *Eocapnosis* | *Eucapnopsis bulba* | EB4 |
| ZOTU04 | Plecoptera | Perlidae | *Kamimuria* | *Kamimuria tibialis* | KT2 |
| ZOTU05 | Plecoptera | Perlidae | *Kamimuria* | *Kamimuria tibialis* | KT5 |
| ZOTU06 | Ephemeroptera | Heptageniidae | *Epeorus* | *Epeorus latifolium* | EL5 |
| ZOTU07 | Ephemeroptera | Heptageniidae | *Epeorus* | *Epeorus latifolium* | EL4 |
| ZOTU08 | Plecoptera | Capniidae | *Eocapnosis* | *Eucapnopsis bulba* | EB3 |
| ZOTU09 | Plecoptera | Perlidae | *Kamimuria* | *Kamimuria tibialis* | KT4 |
| ZOTU10 | No Match | No Match | No Match | No Match | No Match |
| ZOTU11 | Plecoptera | Perlidae | *Kamimuria* | *Kamimuria tibialis* | KT6 |
| ZOTU12 | Ephemeroptera | Heptageniidae | *Epeorus* | *Epeorus latifolium* | EL2 |
| ZOTU13 | Ephemeroptera | Heptageniidae | *Epeorus* | *Epeorus latifolium* | EL3 |
| ZOTU14 | No Match | No Match | No Match | No Match | No Match |
